# Chemical inactivation of two non-enveloped viruses follows distinct molecular pathways

**DOI:** 10.1101/2024.07.16.603687

**Authors:** Pankhuri Narula, Milan Kumar Lokshman, Sandip B. Pathak, Sayandip Mukherjee, Manidipa Banerjee

## Abstract

Non-enveloped viruses, which lack a lipid envelope, typically display higher resistance to disinfectants, soaps and sanitizers compared to enveloped viruses. The capsids of these viruses are highly stable and symmetric protein shells that resist inactivation by commonly employed virucidal agents. This group of viruses include highly transmissible human pathogens such as Rotavirus, Poliovirus, Foot and Mouth Disease Virus, Norovirus and Adenovirus; thus, devising appropriate strategies for chemical disinfection is essential. We tested a mild combination of a denaturant, alcohol, and organic acid on two representative non-enveloped viruses – Human Adenovirus 5 (HAdV5) and Feline Calicivirus (FCV)– and evaluated the molecular pathway of capsid neutralization using biophysical methods. The transition temperatures signifying conformational shifts in the capsid were established in the presence and absence of chemical treatment using Differential Scanning Calorimetry (DSC), while the corresponding morphological alterations were visualized and correlated using Transmission Electron Microscopy (TEM). We found that while chemical treatment of purified HAdV5 particles resulted in increased thermal instability, followed by large scale particle aggregation; similar treatment of FCV particles resulted in complete collapse of the capsids. The distinct effects of the chemical treatment on the morphology of HAdV5 and FCV suggests that non-enveloped viruses with icosahedral geometry can follow different molecular pathways to inactivation. Further, while individual components of the chemical formulation caused significant damage to the capsids, a synergistic action of the whole formulation was evident against both non-enveloped viruses tested. Molecular level understanding of inactivation pathways may result in the design and development of effective mass-market formulations for rapid neutralization of non-enveloped viruses.

**Highlights:**

a. formulation consisting of 3.2% citric acid, 1% urea in 70% ethanol, pH4 effectively inactivates HAdV5 and FCV.
b. inactivation pathways with complete formulation, are different for the two viruses.
c. effect of whole formulation is more effective compared to individual components.

## 1. Introduction

Non-pharmaceutical interventions (NPIs) enforced during COVID-19 had a significant impact on the circulation, transmission and seasonal outbreaks of many other endemic respiratory viruses such as Influenza A virus and Respiratory syncytial virus(1). While the NPIs practiced during pandemic were primarily targeted to reduce airborne transmission of enveloped viruses, very few studies looked at the effect of these measures on the circulation of enteric viruses which are primarily driven by non-enveloped viruses and which transmit primarily through fomites and a fecal-oral route. In a recently published study by Perofsky et al., it was remarkable to note that the viruses that rebounded soon after lifting of restrictions were non-enveloped viruses (Adenovirus, Rhinovirus, and Enterovirus)(2). Non-enveloped viruses are less susceptible to inactivation by lipophilic agents such as surfactants and alcohols(3), demonstrate ability to persist longer on surfaces(4), and transmit significantly better between surfaces upon contact(5). Considering all the above factors, it is likely that non-enveloped viruses continued to circulate and survive on surfaces at levels higher than enveloped viruses, and the removal of pandemic restrictions facilitated an early rebound based on the pre-existing prevalence and low infectious dose required for a productive infection(6). Although enveloped respiratory viruses with a RNA genome have been hypothesized to be of enhanced pandemic potential, the disease burden from periodic outbreaks of diarrhoea and severe gastroenteritis caused by non-enveloped viruses especially in low-resource settings cannot be ignored(7). A global push to develop broadly applicable, instantly deployable antiviral and virucidal strategies is critical to combat challenges posed by existing and emerging infectious viruses. Both hand washing with soap and water, and use of alcohol-based sanitizers, are effective in reducing viral spread when used correctly(8). A hierarchy of pathogens on the basis of susceptibility to antimicrobial chemicals, proposed by E H Spaulding originally in 1957, has placed non-enveloped viruses at a higher level compared to enveloped viruses(9)(10). Only the agents showing substantial inactivation with both group of viruses can be truly categorized as broad-spectrum antiviral agents.

Developing formulations which are effective against non-enveloped viruses, while being safe for human usage, presents unique challenges. Substantial differences have been noted in the degree of inactivation of similar groups of non-enveloped viruses with the same chemical formulation. Interestingly, prominent differences are seen to exist within members of the same genus. In the hierarchy, large non-enveloped viruses like Adenovirus have been placed at a lower level compared to smaller viruses like Noroviruses and Picornaviruses, indicating higher susceptibility to inactivating agents(11). However, within the Adenovirus family, adenovirus 5 shows more susceptibility to ethanol based inactivation, compared to adenovirueses 2 and 8 (12)(3)(13). Also, adenoviruses 2 and 8, in spite of being substantially larger, are less susceptible to ethanol compared to the smaller murine norovirus(14). Several such studies, involving inactivation of non-enveloped pathogens by a variety of chemical agents such as ethanol, isopropanol, sodium hypochlorite, hydrogen peroxide, acid, base, denaturants, quarternery ammonium compounds etc, provide conflicting evidence with regard to the hierarchial classification of viral pathogens on the basis of stability and susceptibility. It has been proposed that the degree of susceptibility to chemical inactivation, specifically for non-enveloped viral pathogens, is primarily due to the chemical nature of the capsid protein coat, which varies from virus to virus, and cannot be normalized on the basis of primary determinants such as size and genome content. For example, a primary determinant of alcohol-based inactivation may be the hydrophobicity of the capsid coat(11). Poliovirus, which has a hydrophilic capsid, is more susceptible to the hydrophilic ethanol compared to isopropanol; while Simian Virus 40 (SV40), with an overall hydrophobic outer surface is more susceptible to isopropanol. Similarly, while Enterovirus 71 and D68 belong to the same genus and have the same size and overall structure, EV-D68 is more acid labile due to its acid-dependent uncoating mechanism(15). Thus, it is clear that a deeper understanding of the physical and chemical pathways of non-enveloped virus inactivation is needed in order to devise broadly active formulations.

This study aims to investigate the mechanism of action of ethanol based chemical formulation against two typical non-enveloped viruses – Human Adenovirus type-5 (HAdV-5) and Feline calicivirus (FCV). HAdV5 is a large DNA virus with a diameter of 80-100 nm, and causes various disorders including respiratory tract infections, conjunctivitis and pharyngitis. Currently there are no approved antiviral therapies against AdVs(16). FCV and Human Norovirus (HNoV) belong to same virus family - *Caliciviridae* - and are comparable in terms of size and structure, which makes FCV the most widely used surrogate system for the study of HNoV(17). It has been established in literature and by global regulatory bodies(18) that the sensitivities of non-enveloped viruses towards inactivating agents can be assessed by studying their activity against these representative particles. In this study, we have utilized a combination of biochemical and biophysical methods to dissect the mechanism of inactivation of HAdV5 and FCV by specific formulations. Our studies show that capsid inactivation mechanism of these representative non-enveloped viruses by the same formulation may vary significantly highlighting the necessity of understanding the molecular inactivation pathways to generate broadly effective virucidals.

## 2. Methods and Materials

### 2.1 Cells and Viruses

The following viruses and cell lines were obtained commercially from the American Type Culture Collection (ATCC), Manassas, Virginia, USA._Human adenovirus type 5, strain Adenoid 75 (ATCC catalogue number VR-5); Feline calicivirus strain F9 (ATCC catalogue number VR-782); HeLa cells (ATCC catalogue number CCL-2). CRFK cells were sourced from the NCCS repository, Pune, India. Cells were maintained in MEM Eagle (Sigma-Aldrich) containing 10% FBS and 1% Pen-strep (Gibco). Viruses were propagated using recommended protocols.

### 2.2 Test formulations

The formulations were made in-lab using commercially available chemicals. The formulation consisted of 3.2% citric acid (Sigma catalogue no. C1909), 1% urea (Merck catalogue no. 0001032342) and 70% ethanol (Merc catalogue number 1009831000) at pH4. The pH was adjusted with 10% triethanolamine (TEA, Sigma catalogue number 90279). A list of tested formulations is presented in Table 2.

### 2.3 Virucidal efficacy of test formulation

Virucidal efficacy tests were conducted using the EN14476 protocol under clean conditions. Briefly, 8 parts sanitizer formulation, 1part 0.3g/L bovine serum albumin (interfering substance) and 1 part virus stock suspension were mixed at room temperature, vortexed and incubated for 1 min. The reaction was immediately stopped by adding 1part Dey-Engley neutralizing broth. 100µl of neutralized suspension was passed through MicroSpin S-400 spin columns at 750g for 2 min at 4°C. The column filtrate was serially diluted in 2% MEM, and 100µl from each dilution was added on to a monolayer of respective cell lines in a 96-well plate. Each dilution was plated in 6-8 wells. The respective virus, neutralization and cell cytotoxicity controls were performed as per the protocol. The plate was then incubated at 37°C in 5% CO_2_, for 5-7 days. The Cytopathic Effect (CPE) observations were then converted to TCID_50_ using the Spearmann-Karber method and log TCID_50_ values were calculated.

### 2.4 Virus Purification

#### 2.4.1 HAdV5 purification

HAdV5 was purified from HeLa cells according to the protocol published by Dumitrescu *et al* (19). Virus concentration was determined using corrected A260 (A260-A320) as described by Maizel *et al*. According to this method, an absorbance of 1.00 AU (with a 1 cm path length) at 260 nm equates 1.1 x 10^12 viral particles per millilitre (20). Purified particles were visualized by negative stain electron microscopy.

#### 2.4.2 FCV purification

CRFK cells were infected with FCV stock at an MOI of 1.0, and virus was purified according to the protocol described by Conley *et al.* (21). Protein concentration of purified particles was estimated using from absorbance at 280 nm (A_280_).

### 2.5 Treatment with chemical formulations

Purified FCV and HAdV5 were treated with the formulation and its individual components, listed in Table 2, for an incubation period of 60 seconds.

### 2.6 Differential scanning calorimetry (DSC)

DSC of untreated and formulation treated virus particles was carried out in respective buffers at a protein concentration of 1 mg/ml (HAdV5) and 1.8 mg/ml (FCV) in a Malvern MicroCal PEAQ-DSC, from 20 to 100°C. In each case, buffer runs were used as sample references. HAdV5 was scanned at a rate of 1°C/min and FCV was scanned at 1.5°C/min.

### 2.7 Transmission electron microscopy (TEM)

4-5 μl of purified virus (HAdV5 and FCV) sample (untreated/ treated) was added to glow discharged grids and allowed to adsorb for 2 mins. The excess sample was blotted with Whatman filter paper and the grid was washed with water thrice and dried. In case of HAdV5, grids were stained twice with 4 μl of 2% uranyl acetate solution, for incubation durations of 10 seconds and 30 seconds, followed by blotting and air drying. For FCV, grids were washed with 2% uranyl acetate (20μl) twice and dried. Finally, 20 μl of 2% uranyl acetate solution was added onto the grids for an incubation time of 5 mins, followed by blotting and air drying. Grids were viewed in a FEI TECNAI TF20 microscope operating at 200 keV, at magnifications ranging from 19000X to 80,000X.

## 3. Results

### 3.1 Virucidal efficacy of alcohol-based formulation containing citric acid and urea

It has been previously reported that citric acid and its derivatives can enhance the virucidal ability of alcohol-based formulations against a range of non-enveloped viruses(14)(22)(23). We therefore evaluated the virucidal ability of alcohol-based formulations containing optimized levels of citric acid and urea to inactivate HAdV5 and FCV within 60 seconds of contact time using a quantitative suspension assay (EN14476) based on European guidelines (18). As shown in table 1, treatment with 70% alcohol alone caused between 1.0 and 1.5 log reduction of input titre for both viruses while the test formulation comprising 70% alcohol along with citric acid and urea showed greater than 4 log reduction within 60 seconds contact time. All the treatments were effectively neutralized by D/E broth. The limit of detection of assay was 2.8 Log TCID_50_ ml-1.

**Table 1.** Virucidal efficacy of test formulations against Human Adenovirus and Feline Calicivirus as determined through a quantitative suspension assay.

| Virus | Test Products | Average Initial viral load<br>(Log<sub>10</sub> TCID<sub>50</sub>/ml) | Average Residual Viral Load<br>(Log<sub>10</sub> TCID<sub>50</sub>/ml) | Average Log10 reduction |
| --- | --- | --- | --- | --- |
| Adenovirus | 70% Ethanol | 6.8± 0.28 | 5.4± 0.14 | 1.4 |
|  | 70% Ethanol+3.2% Citric acid+1% Urea |  | 2.8 | >4.0 |
| Feline Calicivirus | 70% Ethanol | 7.4± 0.14 | 6.6± 0.28 | 1.2 |
|  | 70% Ethanol+3.2% Citric acid+1% Urea |  | 3.0± 0.28 | 4.4 |

### 3.2 Thermal denaturation profiling of HAdV5 and FCV

The effect of the virucidals was tested on the morphology of purified virus particles. The viruses were prepared as per published methods and were found to have expected purification profiles and morphologies (Figure 1, Supplementary Figure1). DSC, a thermoanalytical technique, was utilized to establish a thermal denaturation profile for both viruses. For untreated HAdV5 particles, four distinct transition peaks were noted at 49.1°C, 68.5°C, 71.9°C and 79.2°C (designated T1-T4), indicating sequential conformational alterations (Figure2A). The temperatures align with the reported values for HAdV-5 (Ad5GL variant) (24). For FCV particles, incremental heating resulted in three major transition peaks, with a minor peak at 56.1°C (T1) followed by two major peaks at 62.29°C (T2) and 80.56°C (T3) (Figure 2B). Although there are no prior reports of thermal denaturation studies on FCV, these transition temperatures are consistent with that of other non-enveloped particles of similar size and geometry (25). Higher number of transition states in case of HAdV5 is likely due to more complex, multilayered capsid structure compared to the relatively simpler architecture of FCV.

**Figure 1:**
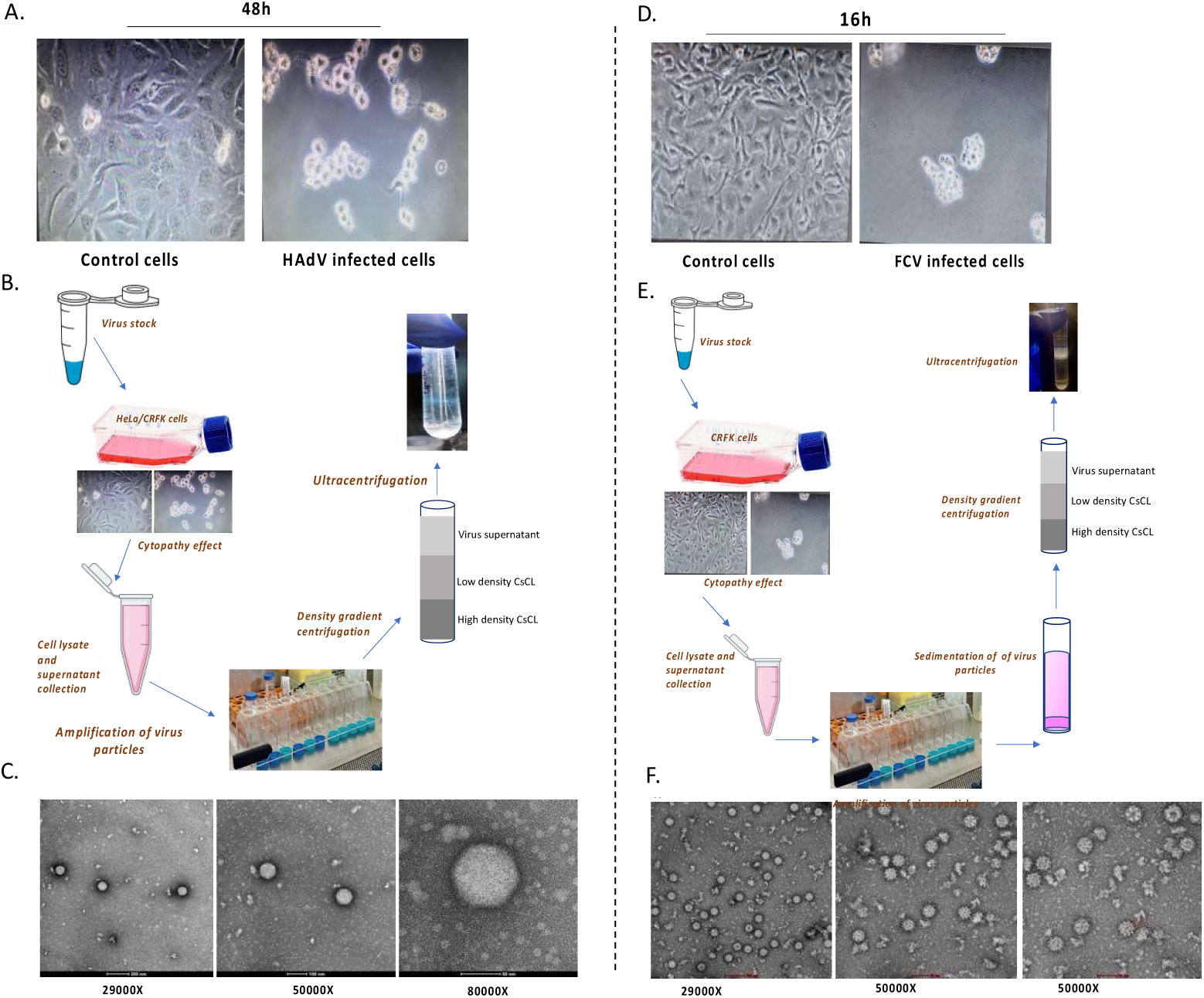
Purification and characterization of non-enveloped viruses. Left panel: Purification and characterization of HAdV5 particles, Right panel: Purification and characterization of FCV particles.

**Figure 2:**
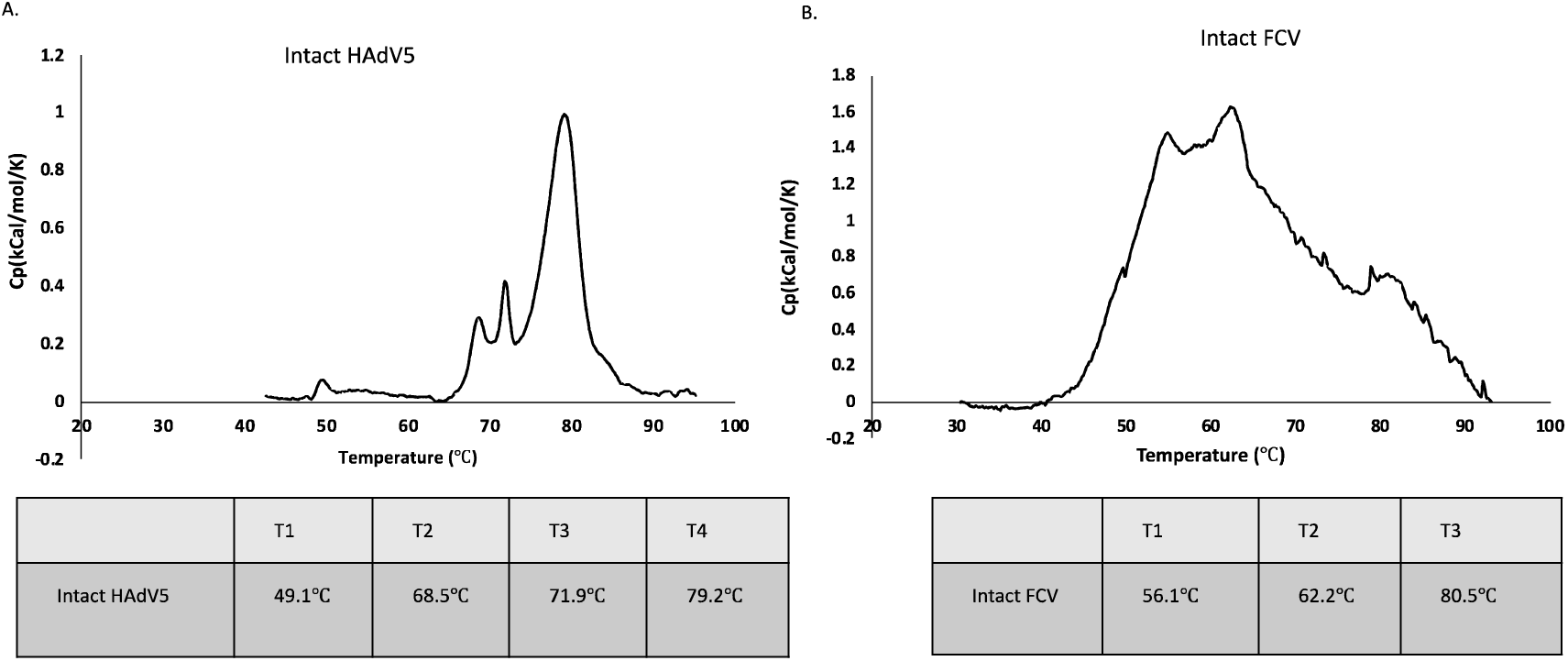
DSC profile of virus particles. (A) Intact HAdV5 and (B) Intact FCV particles were thermally denatured, and the transitions were observed at respective temperatures.

### 3.3 Effect of chemical denaturation on HAdV5 morphology

Purified virus was combined in a 1:1 ratio with four different test solutions-alcohol, urea, citric acid and the complete formulation (Table 2) for 60 seconds, and analysed by DSC and TEM. The DSC profile was expected to provide a qualitative estimate of capsid stability upon treatment, while TEM was employed to directly visualize morphological alterations. We found that the thermal transition of the formulation-treated HAdV5 particles shifted to lower temperatures compared to intact particles (Figure 3A), indicating decreased capsid stability. TEM analysis showed that post-treatment, particles were in a aggregated state with deformed capsids, indicating varying degrees of damage (Figure 3A).

**Figure 3:**
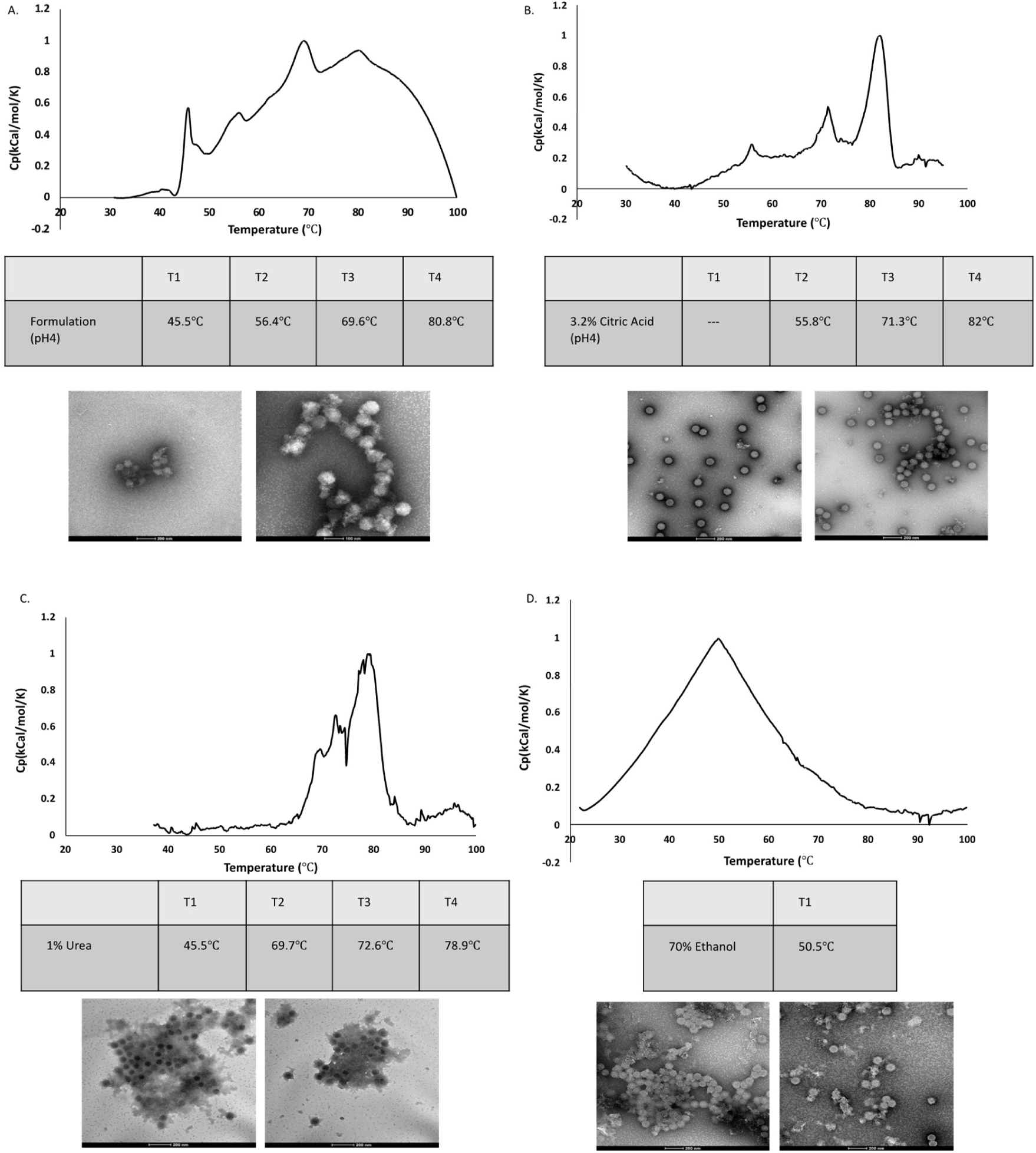
DSC profile and corresponding TEM imaging of HAdV5 particles after treatment with the formulation and its individual components A) treatment with entire formulation consisting of 3.2% citric acid, 1% urea in 70% ethanol (pH4), B) treatment with 3.2% citric acid, C) treatment with 1% urea, D) treatment with 70% ethanol.

**Table 2.** List of test solutions employed for evaluating capsid disruption of Human Adenovirus and Feline calicivirus through biochemical assays and high-resolution imaging.

| Serial number of test products | Composition |
| --- | --- |
| 1. | 3.2% citric acid and 1% urea in 70% ethanol (pH4) |
| 2. | 3.2% citric acid |
| 3. | 1% urea |
| 4. | 70% ethanol |

To understand the role of formulation components on capsid destabilization, HAdV5 particles were treated with individual components. Interestingly, the thermogram of particles treated with 3.2% citric acid at pH 4.0, differed significantly from that produced by formulation treated particles. While no major transition at T1 was observed, T3 and T4 increased (Figure 3B). Concurrent TEM analysis revealed that most virus particles retained structural integrity post-treatment but some aggregated material could be visualized, indicating some capsid component loss (Figure 3B). In contrast, treatment with 1% urea (pH5.3) resulted in a different profile compared to citric acid treatment. While, the first transition T1 was similar to that of formulation-treated particles, but subsequent transitions occurred at higher temperatures. TEM indicated that the majority of the particles were stain filled and highly aggregated (Figure 3C). Treatment with 70% ethanol produced a single major DSC peak at at 50.5°C. Concurrent TEM analysis found that most particles exhibited deformed capsids and were highly aggregated (Figure 3D).

### 3.4 Effect of chemical denaturation on FCV morphology

DSC analysis of the treated FCV particles with the formulation resulted in a single, broad transition peak between 55-65°C (Figure 4A). Concurrently, TEM analysis showed completely dissociated particles, indicating significant disruption in capsid structure (Figure 4A). This result differed from that with HAdV5, where similar transition peaks were observed in the DSC pre- and post-treatment; albeit they shifted to lower temperatures indicating a decrease in stability. The effect of individual formulation components on FCV morphology varied significantly from that of the whole formulation. Treatment with 70% ethanol produced two transition peaks at 48.61°C, and 63.01°C in the DSC thermogram (Figure 4B), suggesting sequential disassembly events rather than complete rupture. TEM analysis of the ethanol treated samples showed partially damaged as well as intact particles, unlike the damaged ones observed with the whole formulation (Figure 4B). Treatment with ethanol also resulted in the production of smaller, homogeneous assemblies indicative of T=1 particles or capsomeres, suggesting an entirely different dissociation pathway compared to the whole formulation.

**Figure 4:**
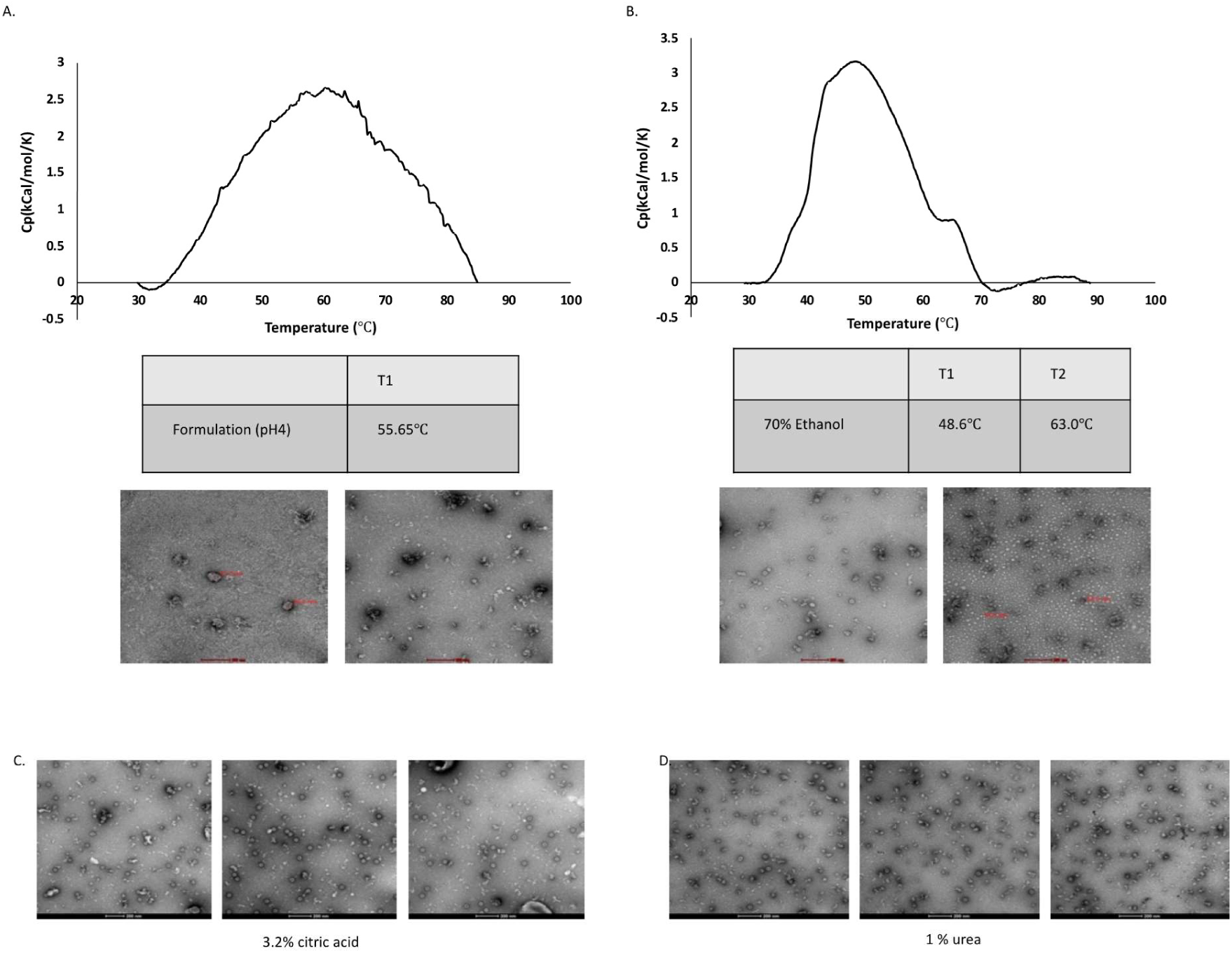
Thermal and morphological characterization of FCV particles after treatment with formulation. a) treatment with entire formulation, b) treatment with 70% ethanol alone c) treatment with 3.2% citric acid, d) treatment with 1% urea.

Treatment with 3.2% citric acid predominantly preserved the FCV structural integrity, maintaining the characteristic cup-shaped appearance (Figure 4C). In contrast, 1% urea treatment resulted in mixed population of partially and completely damaged particles along with some intact ones. (Figure 4D).

### 3.5 Quantification of morphological damage

In order to quantify the effect of the complete formulation, and its individual components, on HAdV5 and FCV, visual classification of the damaged particles were carried out from negatively stained micrographs. For this purpose, a total of 100 random particle images were considered from each group - a) complete formulation, b) 3.2% citric acid, c) 1% urea and d) 70% ethanol. The particles were visually categorized into: a) intact, b) semi-damaged: exhibiting visible capsid damage, while retaining spherical shape and c) damaged: completely collapsed or aggregated particles with no defined shape and size. For both viruses, the maximum percentage of damaged particles was observed with the complete formulation. In case of HAdV5, 70% ethanol caused the maximum disruption among all components, generating 78% damaged particles. Treatment with 3.2% citric acid and 1% urea resulted in higher percentages of semi-damaged particles (87% and 86% respectively), with negligible particles in damaged group (Figure 5A).

**Figure 5:**
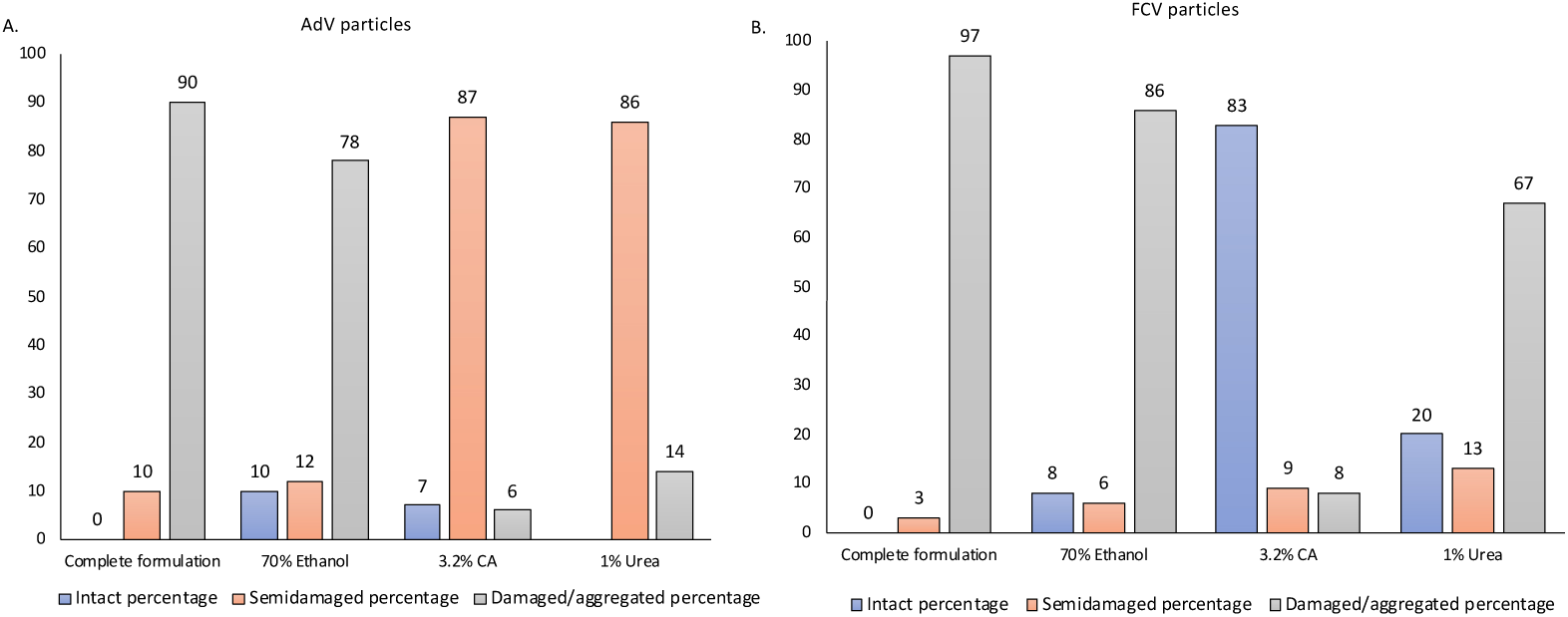
Quantification of HAdV5 and FCV particles, with distinct morphologies, from TEM micrographs.

In case of FCV, ethanol and urea caused the maximum damage, with 86% and 67% of the particles categorized as damaged respectively. Citric acid alone caused almost negligible damage. However, the majority of the damaged particles (97%) were observed with the complete formulation (Figure 5B).

Our data indicates synergistic effect of the formulation components on particle morphology in both viruses, with no individual component matching the level of disruption achieved by the entire formulation.

## 4. Discussion

The pandemic potential of non-enveloped viruses, that resist normal hand hygiene modes, highlights the urgent need for suitable inactivation strategies as well as molecular studies on the mechanism of inactivation. While the lability of the outer lipid coat to soaps and alcohol-based sanitizers makes systematic inactivation of enveloped viruses possible; most non-enveloped viral pathogens are difficult to neutralize within a short contact time of 30-60 s. Antiviral agents like bleach can dissociate non-enveloped virus capsids but are unsuitable for human use (26). No current hand hygiene formulations are effective against both enveloped and non-enveloped viruses..

We formulated an effective neutralizer for non-enveloped viruses combining 3.2% citric acid with 1% urea in a base of 70%. ethanol. We tested it against two representative viruses – HAdV5 and FCV, a surrogate for HNoV, both with icosahedral geometry. The test formulation was effective in causing four log decrease in the titers of both viruses within a minute of contact in a standardized suspension assay. Given the lack of mechanistic studies on non-enveloped virus capsid inactivation by chemical agents, we further attempted to investigate the effect of the formulation and its components by assessing the thermal denaturation profile of viruses using DSC, followed up with TEM for morphological studies.

It has been established by our laboratory, as well as others, that this energy landscape during gradual disassembly of the capsids can be captured *in vitro* by incremental heating of the purified capsids (25). We conjectured that formulation-treated capsids would exhibit distinct thermal denaturation profiles compared to untreated ones, reflecting the nature of inflicted damage. While the formulation effectively neutralized both HAdV5 and FCV, the latter being a highly resistant norovirus surrogate, however, the mechanisms of inactivation differed. While HAdV5 particles underwent stepwise loss of crucial elements such as the capsid fibres, capsid proteins and eventually aggregated; FCV particles were entirely dissociated into smaller complexes. While formulation-treated HAdV5 still displayed stepwise transitions, although with decreased peak temperatures indicating thermal instability; treating FCV with the formulation completely altered the thermal denaturation profile, resulting in a broad single peak instead of three major transitions seen in untreated particles. In order to comprehend the variance in inactivation mechanisms, we investigated the surface chemistry of the viruses (Figure S2). Both HAdV5 and FCV displayed similar surface chemistry characteristics, suggesting that the variation in their inactivation response to the treatment formulation is not solely due to surface charge differences. Furthermore, our results do not adhere to the conventional Spaulding classification, which typically associates higher resistance with small non-enveloped viruses.

It was also clear that the complete formulation was more effective than the individual components in hastening disassembly or causing dissociation of virus components. It is well established that 70% ethanol, the primary constituent of commonly available sanitizers, can exert a certain degree of damage to the virus capsid. The inactivating effect of ethanol on adenoviruses has been noted at concentrations ranges of 50-70% (27). A previous study has reported that 70-90% ethanol (at its own pH) is effective against HAdV5 at a treatment time of 30-60 seconds(3). Our results show that while the treatment of HAdV5 with 70% ethanol resulted in significant capsid disruption and the aggregation of virus particles, the entire formulation caused marginally more damage. However, application of 70% ethanol to FCV resulted in a substantial proportion of virus particles remaining intact. Particularly noteworthy was the presence of T=1 viral particles. Thus, 70% ethanol alone may not be sufficient to inactivate non-enveloped viruses, although it does cause some damage.

The greater effect of alcohol on HAdV5 is in agreement with the Spaulding hierarchical classification, which suggests that large non-enveloped viruses are more susceptible to inactivating agents like alcohol compared to relatively smaller ones like NoV. A few other studies have attempted to combine other denaturants including urea, citric acid and 2-propanol with ethanol (28)(23). However, in our study, the constituents of the formulation other than 70% ethanol exhibited varying effects of comparatively lower magnitude when individually used. The DSC thermogram of HAdV5 treated solely with 3.2% citric acid, showed higher transition temperatures compared to particles treated with the complete formulation, indicating that the formulation elicits greater degree of capsid instability compared to citric acid alone. Also, citric acid alone exhibited relatively limited efficacy in FCV inactivation, as a substantial proportion of intact particles were observed post-treatment. While the denaturing property of urea resulted in reduction in capsid integrity in both HAdV5 and FCV, the overall effect was less than that observed with the complete formulation. Previous studies on urea-based virus inactivation support these findings (29)(30).

In summary, our combined thermal and morphological analysis suggests that the components of our formulation function synergistically to affect the stability of the non-enveloped virus particles. While each individual component has measurable but distinct effect on particle stability, a more profound cooperative effect on virus inactivation is observed with the complete formulation. Surprisingly, while both HAdV5 and FCV are icosahedral non-enveloped particles, the mechanistic pathways to inactivation appear distinct (Figure 6). A better understanding of particle chemistry and mechanism of inactivation is needed in order to devise broadly effective denaturing agents.

**Figure 6:**
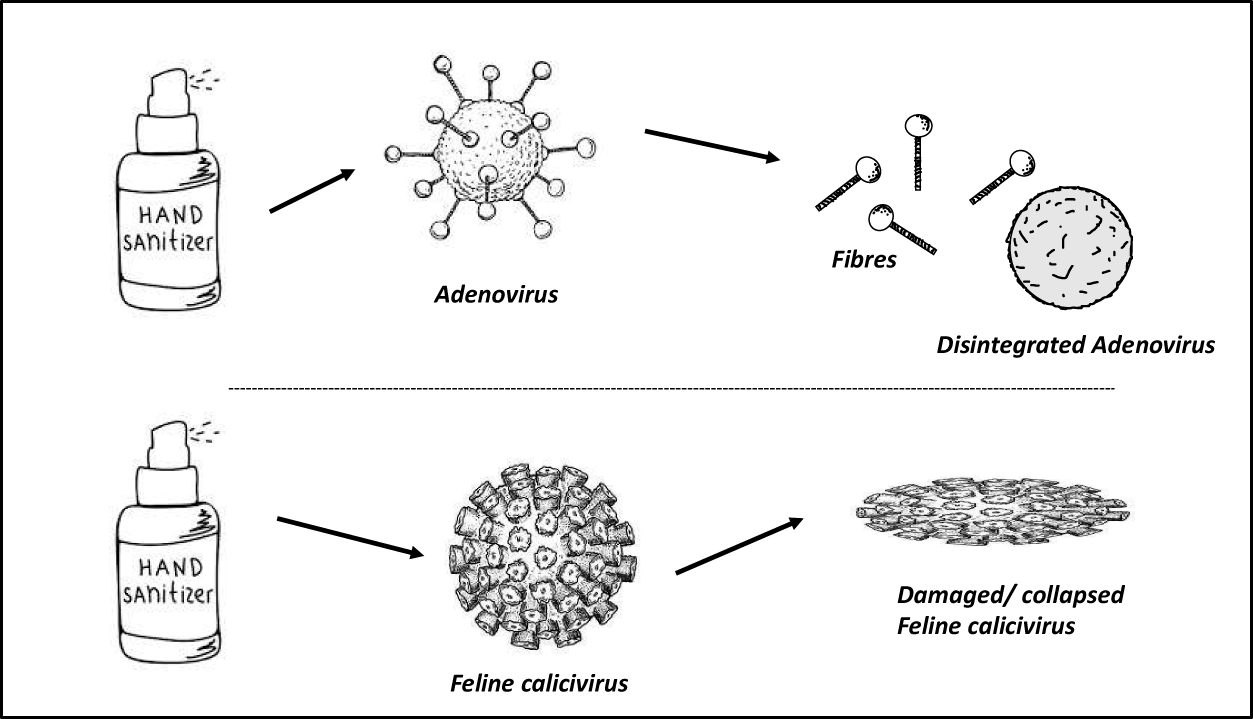
Schematic representation of the mechanism of inactivation of HAdV5 and FCV after treatment with complete formulation pH4.

## Acknowledgements

This work is supported by Unilever, Grant number FT/5/288/2020 given to MB. The authors thank the Central Research Facility IIT Delhi, and Akshey Kaushal, for help with TEM imaging; and IIT Delhi, Grant number PLN03/EBSC, Prof. Bishwajit Kundu, and Dr. Shubham Vashishtha for help with DSC data acquisition and analysis. PN thanks the Department of Biotechnology (DBT) for grant (BT/PR50894/BIC/101/1317/2023) under the DBT-BioCARe scheme, and MKL thanks DBT for fellowship support.

## Author contributions

Conceptualization: MB, SM, Methodology: PN, MKL, SP, SM, MB; Investigations: PN, MKL, SP, Writing-original draft: PN, MKL, SP; Writing, review and editing: PN, MKL, SP, SM, MB; Supervision; MB, SM; Project administration; SM, MB.

## Abbrevations

ATCC: American Type Culture Collection
CsCl: Cesium Chloride
CPE: Cytopathic Effect
DSC: Diffrential Scanning Calorimetry
FCV: Feline Calicivirus
HAdV: Human Adenovirus type-5
HNoV: Human Norovirus
MOI: Multiplicity of Infection
NCCS: National Centre for Cell Science
SV40: Simian Virus 40
TCID50: 50% Tissue Culture Infection Dose 12.TEA-Triethanolamine
TEM: Transmission Electron Microscopy

