## Supplementary Material for "Chemical inactivation of two non-enveloped viruses follows distinct molecular pathways"

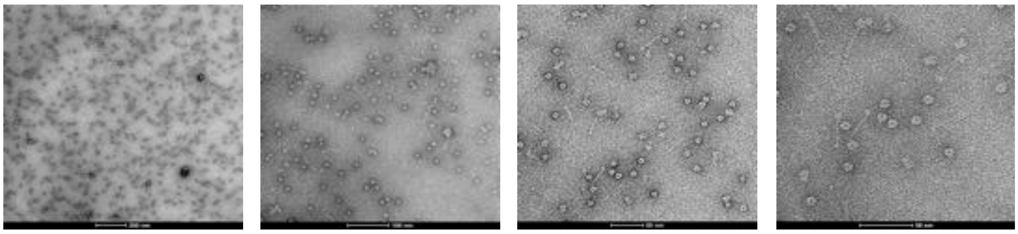

Figure S1: Morphological characterization of HAdV5 purified from upper band after CsCl density centrifugation

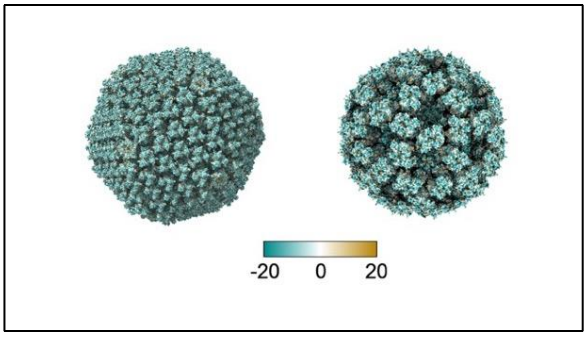


Figure S2: Surface hydrophobicity distribution in a) HAdV5 and b) FCV.
